# Does sample size of leaf osmotic potential affect its relationship with cotton yield?

**DOI:** 10.1101/2025.01.17.633093

**Authors:** Xuejun Dong, Dale A. Mott, Quan Zhou, Benjamin M. McKnight

**Affiliations:** Texas A&M AgriLife Research, Uvalde, Texas 78801, USA; Texas A&M AgriLife Extension Service, College Station, Texas 77843, USA; Department of Statistics, Texas A&M University, College Station, Texas 77843, USA

**Keywords:** cotton, leaf osmotic potential, sample size

## Abstract

Leaf osmotic potential at full turgor (*π*_0_) has been used frequently to indicate turgor loss point of plant leaves. However, even a rapid measurement of *π*_0_ using osmometry is time-consuming, if numerous leaf samples need to be measured. Because of this, researchers tend to use a small sample size to determine *π*_0_ and relate it to indices of crop performance. Yet the statistical and agronomic significance of using a small sample size of *π*_0_ to indicate crop performance is not known. We address this question using field measurements and statistical resampling. Six mature leaf samples were collected at the peak bloom stage from each of the 54 cotton plots in Texas, USA in 2024. The *π*_0_ of the collected leaves were measured using an osmometer. Seed cotton yields from the field plots were measured near the end of cotton season. To test the effect of sample size on strength of the linear relation between *π*_0_ and cotton yield, 1-6 resamples of *π*_0_ were randomly drawn with replacement from the original 6 measurements per plot for the 54 plots. The resampled data of *π*_0_ were then used as independent variable to predict cotton yield. We found that, considering the labor and cost, sampling 3 or 6 leaves per plot may not make a significant difference for the linear regression between *π*_0_ and cotton yield.

## Introduction

Leaf osmotic potential at full turgor (*π*_0_) has been shown to indicate leaf turgor loss point, an important index of plant drought tolerance (1). The reason for choosing to measure leaf osmotic potential at full turgor, but not at an arbitrary water content, is to minimize the effect of leaf water content on the measured value of osmotic potential (2). This parameter was previously determined through the construction of the so-called pressure-volume (PV) curve, in which *π*_0_ was estimated through the analysis of the data of leaf water potential and leaf water content collected from various time points during the dehydration process of a sample leaf, starting from the state of full saturation (3). It typically required more than 10 hours to build a single PV curve. Recently, Bartlett et al. (1) proposed to use osmometry to rapidly determine *π*_0_, leading to 60-fold increase in measurement speed, when compared with the traditional PV method. In recent decade, this new method has been adopted by numerous researchers (4–11). However, since it may take 15-20 minutes to measure one sample using this rapid method, it is still time-consuming, if numerous leaf samples need to be measured for drought tolerance survey. Dong et al. (12), when testing drought tolerance capacity of cotton varieties in Texas, used 3 leaf samples per plot to measure *π*_0_ and relate it to the measured cotton yield and fiber quality indices. Yet, it is unknown whether and to what extent using small sample size may affect the regression relations between *π*_0_ and cotton yield. This question is addressed in the current study by combined use of field measurements and statistical resampling.

## Materials and Methods

The study sites are located at Hutto, Texas, USA (30°32’40”N, 97°32’43”W) and Uvalde, Texas, USA (29°12’55”N, 99°46’41”W). The average annual temperatures of Hutto and Uvalde are 20.3 °C and 21.7 °C, respectively, and the amounts of annual rainfall received are 918 mm and 566 mm, respectively. The soil at Hutto site is Branyon clay with 29% sand, 23% silt and 49 % clay. The soil at the Uvalde site is Knippa clay with 23% sand, 30% silt and 47% clay. The Hutto site was not irrigated (dryland) and received 216 mm of rain during the crop season in 2024. The plots at Uvalde site were managed under deficit irrigation, receiving 308 mm of rain plus irrigation during the 2024 cotton season. At the early growth stage (measured on May 3, 2024), the Uvalde site had 50% of available water in top 1-m soil profile. The initial soil moisture at the Hutto site was not measured, but was likely very high due to the abundant rains received weeks prior to planting.

Cotton leaf samples were collected from fields at Hutto, Texas (97-cm row spacing, planted/harvested on May 24/Sep. 30) and Uvalde, Texas (76-cm row spacing, planted/harvested on Apr. 16/Sep. 15) at the peak bloom stage in 2024. Six mature leaves from the 4th node (counting from top) were collected from each of 30 plots at Hutto and 24 plots at Uvalde. The leaves were cut from the petiole base, immediately submerged under distilled water, and brought to a laboratory for *π*_0_ to be measured using a 5520 VAPRO Vapor Pressure Osmometer (Wescor, Inc., USA), following Bartlett et al. (1) and Dong et al. (12). Briefly, the leaves were stored in a bucket over night under high-humidity condition, with the cut end of the petiole submerged in a thin layer of distilled water. The next morning, the water droplets on the leaf surfaces were gently removed by blotting using tissue paper. Then, a 8-mm diameter leaf disc was punched off each of the leaves, wrapped in aluminum foil, and snap-frozen in liquid nitrogen. The frozen samples were then promptly transferred to a -80 °C freezer awaiting measurement using osmometry. The osmometer method of measuring *π*_0_ involved recording a series of successively decreasing values of the osmolality readings over a time period of approximately 15 to 20 minutes, during which the frozen leaf disc underwent the thawing process while being sealed in the sample holder of the osmometer. The process was terminated (i.e., the measurement completed) when either of the following two scenarios occurred: (a) the osmolality reading started to repeat the prior value; and (b) the reading started to increase. This was presumably the point at which the maximum amount of vacuolar liquid was released from the thawing leaf disc and some other artifacts, such as the dissolved solutes from the broken cell wall structures (1), were not yet influencing the vapor pressure in the immediate vicinity of the sealed leaf disc.

The unit of the readings from the osmometer was in mmol/kg, which was converted to the volume-based water potential unit, such as MPa, according to the Van’t Hoff equation (13):

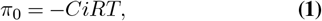

where *C* was the osmolality value in mol/kg, *i* is an ionization constant assumed to equal to unity, *R* is the ideal gas constant (0.0083143 kg MPa/mol/K), and *T* the absolute temperature (*K* = °C +273).

Seed cotton yields from all the 54 field plots were measured near the end of cotton season. At the Hutto site, the whole plots (6 rows *×* 351 m) were harvested using a cotton stripper. At Uvalde, each plot was harvested by hand using 2 rows, 4.6 m segments.

To test the effect of sample size on strength of the linear relation between *π*_0_ and cotton yield, 1-6 resamples of *π*_0_ measurements were randomly drawn (with replacement) from the original 6 measurements per plot for the 54 plots. The average *π*_0_ value of each of the 54 plots was used as a value of independent variable (*x*) to predict cotton yield (*y*). For each regression, the correlation coefficient (*r*) was calculated. Also calculated was the average uncertainty of *y, σ*_*y*_, according to Taylor (14):

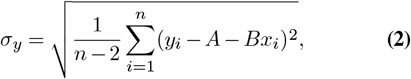

where *n* = 54, and *A* and *B* were calculated respectively as

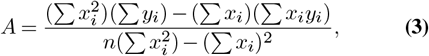

and

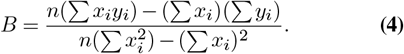

The above process was repeated 200 times, i.e., by drawing 200 sets of averaged *π*_0_ value for each of the 54 plots with a particular sample size of 1, 2, 3, 4, 5 or 6 leaves per plot. A computer program was written to automate the processes of drawing random resamples and conducting regression analysis using the resampled data values of *π*_0_ and the field-measured seed cotton yield. Finally, the resultant values of *r* and *σ*_*y*_ associated with different sample sizes were compared using the One-way ANOVA (*p* = 0.05).

## Results and Discussion

The typical time dynamics of the readings of the osmometer (in mmol/kg) during the thawing process of two example frozen leaf discs are shown in Fig. 1A. The main reason for waiting for 80 s between adjacent readings is to allow thermodynamic equilibrium to be established between the thawing leaf disc and its immediate micro-environment in the sample holder (15). The linear regression between *π*_0_ and cotton yield using the original six measurements of *π*_0_ is shown in Fig. 1B. The histograms (with normal fits) of *r* and *σ*_*y*_ for the linear regressions using 200 sets of 1-6 resamples of *π*_0_ samples for each of the 54 cotton plots are shown in Figs. 2 and 3. It can be seen from these figures that both the *r* and *σ*_*y*_ values were widely dispersed when the sample size of *π*_0_ from each cotton plot was limited to 1-2, indicating the poor strength of the resultant linear regressions. On the other hand, the shapes of the distributions of the *r* and *σ*_*y*_ values, as well as the respective mean values and standard deviations, were similar for the resultant regressions associated with samples sizes of 3-6 for *π*_0_.

**Fig. 1.**
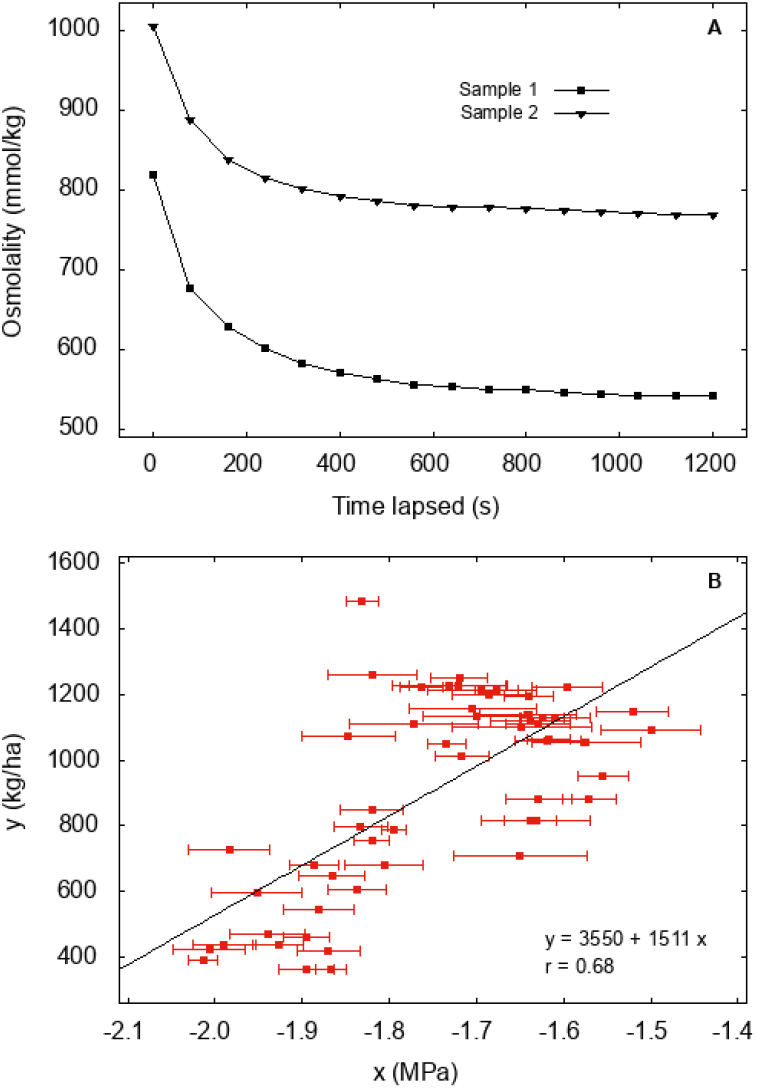
(**A**) Recorded osmolality readings during the thawing process of two random samples of frozen leaf discs. The osmolality reading of the penultimate data point from each series of readings was considered the final osmolality reading of that sample. (**B**) Linear regression of measured seed cotton yield (kg/ha) in relation to average leaf osmotic potential *π*_0_ of 54 cotton plots. Six leaves were measured in each plot and their average was used as *x*-variable representing each data point in the figure. The error bars indicate the *±*1 standard errors of means of *π*_0_. Also shown are the regression equation and the correlation coefficient (*r*).

**Fig. 2.**
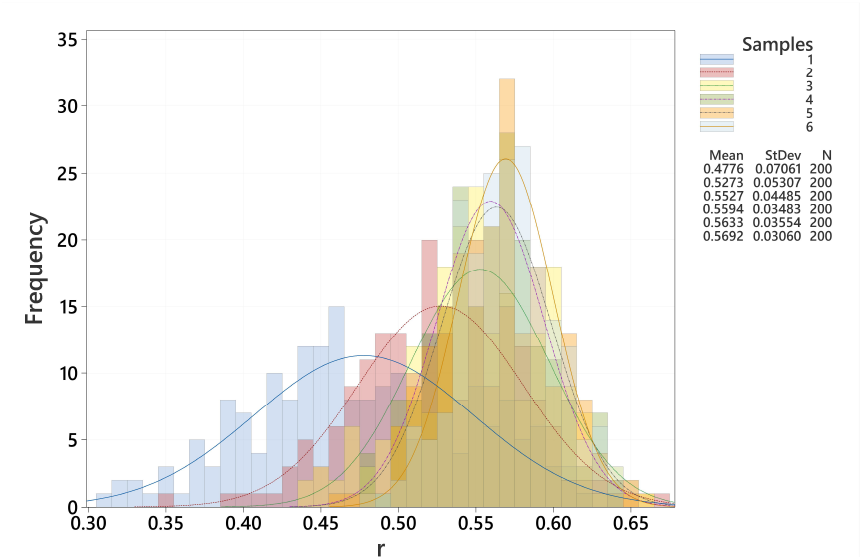
Histograms of *r* of the linear regressions using resampled values of *π*_0_.

**Fig. 3.**
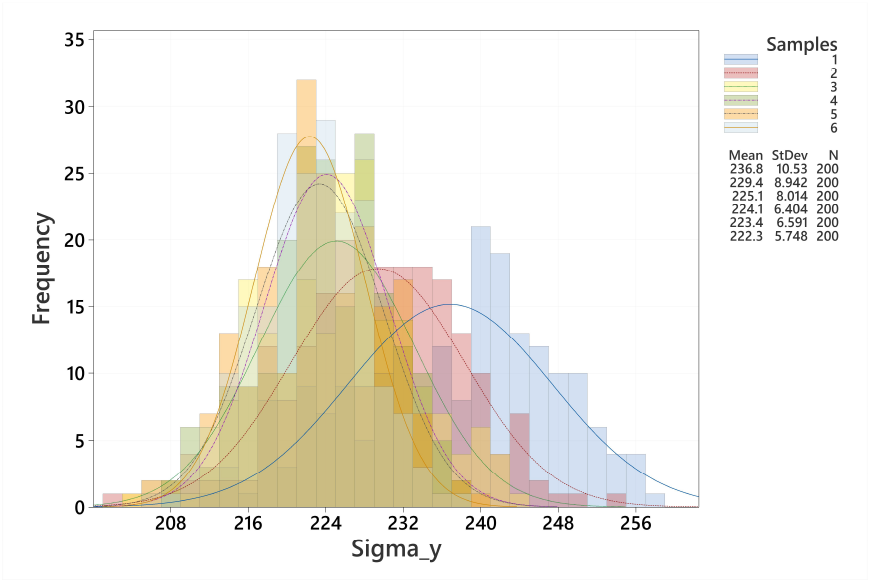
Histograms of *σ*_*y*_ of the linear regressions using resampled values of *π*_0_.

The mean values of *r* and *σ*_*y*_ for the linear regressions using resampled *π*_0_ are compared in Table 1. Our results suggest that increasing sample size led to improved strength of the resultant linear regressions between *π*_0_ and seed cotton yield. However, doubling the sample size from 3 to 6, and thus also of the workload, only reduced the average uncertainty of the predicted cotton yield (*y*) by 1%. As a result, considering the labor and cost, it appears that sampling 3 or 6 leaves per plot may not make a significant difference for the linear regression between leaf osmotic potential and cotton yield. This supports the work of Dong et al. (12), who sampled three leaves per plot to measure *π*_0_ and established a strong relationship between *π*_0_ and selected cotton yield and fiber quality indices for data collected from multiple sites over three years. Our results also suggest that, when a modest sample size of *π*_0_ is acceptable, the researchers’ time/efforts can be invested in collecting data of *π*_0_ for a large number genotypes or treatments to establish a broad relationship between *π*_0_ and cotton yield. Whether the same conclusion holds for other plant species remains to be tested, but we provide the computer code and sample dataset to facilitate the extension of the results of this work to other plant species growing in semi-arid or arid environments where water stress is a driving factor limiting their growth and development.

**Table 1.** Mean values of *r* and *σ*_*y*_ for the linear regressions using resampled *π*_0_.

| Sample size | $r$ | $\sigma_y$ (kg/ha) | N |
| --- | --- | --- | --- |
| 1 | 0.478<sup>D</sup> | 265.5<sup>A</sup> | 200 |
| 2 | 0.527<sup>C</sup> | 257.2<sup>B</sup> | 200 |
| 3 | 0.553<sup>B</sup> | 252.3<sup>C</sup> | 200 |
| 4 | 0.559<sup>AB</sup> | 251.2<sup>CD</sup> | 200 |
| 5 | 0.563<sup>AB</sup> | 250.4<sup>CD</sup> | 200 |
| 6 | 0.569<sup>A</sup> | 249.3<sup>D</sup> | 200 |

## Acknowledgements

Funding support from Cotton Inc./Texas State Support Committee, as well as from USDA-NIFA Hatch project 9574-2, is appreciated. We thank Jose Teran and Joe Gonzalez, Farm Manager and Farm Foreman, respectively, at Uvalde Research Center, and collaborating farmer Rick Kruger for time/efforts invested in crop management.

## Data availability

The data and computer code for reproducing the results of this paper are available from https://zenodo.org/records/14635663.

